# Higher methanotroph abundance and bottom-water methane in ponds with floating photovoltaic arrays

**DOI:** 10.1101/2025.07.24.666521

**Authors:** Nicholas E. Ray, Sophia Aredas, Steven M. Grodsky, Ash Canino, Simone J. Cardoso, Meredith A. Holgerson, Meredith Theus, Marian L. Schmidt

## Abstract

Floating photovoltaic (FPV) arrays alter the methane (CH_4_) cycling dynamics of waterbodies on which they are deployed. Here, we investigated dissolved CH_4_ dynamics and associated CH_4_ cycling microbial communities (methanogens and methanotrophs) in the second year of FPV deployment (70% aerial coverage) in experimental ponds. We found that bottom-water CH_4_ concentrations were twice as high in ponds with FPV compared to those without, while surface water CH_4_ concentrations were orders of magnitude lower than bottom-waters, but did not differ between treatments. There was no change in the relative abundances of putative sediment methanogens or methanotrophs, but FPV restructured methanogen communities. FPV promoted late-summer methanotroph blooms in the water column, with abundances surpassing 1,000,000 cells mL^-1^. We conclude that prolonged periods of CH_4_ production in low oxygen FPV ponds favored blooms of methanotrophs, that may mitigate diffusive CH_4_ emissions to the atmosphere by consuming dissolved CH_4_.

## Introduction

Methane (CH_4_) emissions from aquatic ecosystems have increased because of human activity (Jackson et al. 2024). Floating photovoltaic arrays (FPV), first deployed in 2007, are a rapidly expanding renewable energy technology that can alter aquatic CH_4_ dynamics and emissions (Cazzaniga and Rosa-Clot 2021; Ray et al. 2024). By 2023, over 643 FPV power plants had been installed on lakes, ponds, and reservoirs worldwide (Nobre et al. 2024; Ramanan et al. 2024). There is substantial interest and potential in expanding FPV use on reservoirs and lakes to meet demands for renewable energy (Jin et al. 2023; Woolway et al. 2024), but the ecological and biogeochemical tradeoffs – particularly effects on CH_4_ cycling – must be evaluated before broader deployment (Almeida et al. 2022; Gallaher et al. 2025).

FPV deployment is associated with lower water temperatures (Dörenkämper et al. 2021; Andini et al. 2022), reduced oxygen availability (Wang et al. 2022; Ray et al. 2024), and changes in stratification (Ilgen et al. 2023). These factors influence CH_4_ dynamics in small, shallow systems where FPV has most commonly been deployed (Bastviken et al. 2004; Aben et al. 2017; Ray and Holgerson 2023; Nobre et al. 2024). Indeed, in the first year of FPV deployment, experimental ponds with FPV showed twice the dissolved CH_4_ concentrations and CH_4_ ebullition compared to control ponds, but diffusive CH_4_ emissions were lower (Ray et al. 2024). Whether these observed patterns persist over time or might be similar in other ecosystems is unclear.

Understanding how FPV affects methanogens (CH_4_ producers) and methanotrophs (CH_4_ consumers) is essential for predicting CH_4_ emissions. Methanogenic archaea and methanotrophic archaea and bacteria jointly regulate CH_4_ dynamics, yet it remains unresolved how FPV-induced changes in temperature, light, oxygen, and primary production influences their abundance, composition, and activity. Evidence from non-FPV settings suggests changes in such limnological conditions can re-structure CH4-cycling communities and alter metabolic rates (Wang et al. 2021; Bertolet et al. 2022). Clarifying microbial responses to FPV is critical for determining whether this technology amplifies or dampens CH_4_ emissions and, more broadly, for understanding how pond microbial communities respond to reduced light and oxygen.

Here, we investigate how FPV affects CH_4_ dynamics in temperate experimental ponds during the second year of FPV deployment, with a focus on identifying microbial and biogeochemical mechanisms underlying dissolved CH_4_ concentrations. We pair observations of temperature, dissolved oxygen, and dissolved CH_4_ with measurements of methanogen and methanotroph community structure and CH_4_ process rates during an intensive mid-summer sampling campaign in ponds with and without FPV. To capture microbial mechanisms relevant to dissolved CH_4_ concentrations, we focus on well-characterized methanotrophs and integrate community data with functional rate measurements to capture both composition and activity.

## Methods

### Location and Sampling Scheme

In 2023, FPV arrays (70% aerial coverage) were deployed on three ponds at the Cornell Experimental Ponds Facility in Ithaca, New York, USA (42.5049 N, 76.4666 W). Observational sampling of six ponds starting in 2022 prior to FPV deployment in 2023, providing three replicate control and three replicate FPV ponds (Ray et al. 2024). Each pond is a 30 x 30 m inverted, truncated pyramid, 1.75 m deep, unlined, and with sediment accumulation since construction in 1958-1959. Rooted, submerged macrophytes dominate primary production.

We returned in summer 2024 to continue measurements of dissolved CH_4_ (surface and bottom), temperature, and dissolved oxygen (4 depths; 0.1, 0.75, 1, and 1.25 m from the water surface). We sampled methanogens and methanotrophs four times in 2024 from surface water, bottom water, and sediments. On July 11, 2024, we measured diffusive CH_4_ exchange and net CH_4_ production or consumption in surface waters, bottom waters, and sediments. For each date, we calculated density gradients of pond water columns using temperature profile data to compare mixing and stratification dynamics between ponds with and without FPV (Supplemental Methods).

### Observational CH_4_ Concentration Measurements

We collected samples to determine dissolved CH_4_ concentrations in the surface and bottom water of each pond six times between June 20 and September 5. Samples were collected in triplicate from the pond center using a headspace equilibration approach, with surface samples collected by hand at 10 cm depth and bottom water collected using a Van-Dorn bottle ∼10 cm from the sediment-water interface (Ray et al. 2024). Samples were stored in 12 mL pre-evacuated borosilicate exetainer vials until analysis using a gas chromatograph equipped with a flame ionization detector (Shimadzu GC-2014).

### Microbial Sample Collection and Preservation

Water samples for microbial analysis were collected as for CH_4_ measurements and sequentially filtered through 200 μm and 20 μm NYTEX mesh into 2 L Nalgene bottles and kept on ice until return to the laboratory the same day. Surface sediments (top 2 cm) were collected using a PVC pole corer and stored in WhirlPak bags in the cooler. Water was filtered onto 0.22 μm polyethersulfone filters using a peristaltic pump (190-2020 mL per sample). Sediments were homogenized using a stainless-steel coffee grinder (three 1 second pulses per sample). Filters (halved) and homogenized sediment were stored in 2 mL cryovials at −80 °C. Samples for flow cytometry were fixed with 1 μL 25% glutaraldehyde, diluted 15–20×, stained with SYBR Green I, and run in triplicate on an Attune NxT (*see supplemental methods*).

### Microbial DNA Extraction, Sequencing and Bioinformatics

DNA was extracted from half filters using the DNeasy PowerWater Kit and from sediments using the DNeasy PowerSoil Pro Kit (Qiagen), eluted in 50 μL elution buffer, quantified with a Qubit fluorometer (Thermo-Fisher) and stored at −20 °C. The V4 region of the 16S rRNA gene was amplified following the Earth Microbiome Project protocols (Caporaso et al. 2011; Apprill et al. 2015; Parada et al. 2016) using triplicate 25 μL PCRs with KAPA HiFi 2X Mastermix (Roche), 0.2 μM 515F (5’-GTGYCAGCMGCCGCGGTAA) and 806R (5’-GGACTACNVGGGTWTCTAAT) primers with Illumina Nextera overhangs, and 5 ng template DNA. Zymo-BIOMICS Microbial Community DNA Standard (Zymo) was included to assess error rates. Samples, including PCR and indexing blanks, were pooled, indexed, and sequenced (2 x 250 bp) on an Illumina MiSeq sequencer at the Cornell Biotechnology Resources Center.

Raw sequences were processed into amplicon sequence variants (ASVs) using the DADA2 workflow (Callahan et al. 2016) in R. Reads were trimmed to 253-254 base pairs (max error = 1), merged, denoised, and chimera-checked. Taxonomy was assigned with GreenGenes2 (McDonald et al. 2024). Data were imported into *phyloseq* (McMurdie and Holmes 2013); mitochondrial, chloroplast, and ASVs with a higher relative abundance in blanks were removed. The mock community revealed 24 spurious ASVs (error rate of 0.087%). Samples with low read counts were excluded, resulting in 8,188-171,275 reads per sample. Water column ASV relative abundances were multiplied by cell counts to obtain absolute ASV counts per mL whereas the ASV relative abundances in sediments were multiplied by the minimum library size (20,826 sequences) for relative abundance.

Microbial metabolic profiles were predicted using custom code Functional Annotation of Prokaryotic Taxa version 2 (FAPROTAX v2) based on cultured prokaryotes (Louca et al. 2016). Using default parameters, ASVs were putatively identified through phylogenetic placement onto the FAPROTAX v2 reference tree. Methanogens and methanotrophs were identified at the family and genus level and verified through scientific literature and public databases (NCBI; Table S1).

### Diffusive CH_4_ Flux Measurements and Estimation of k600

We measured diffusive fluxes at the center and edge of each pond using floating chambers (18.93 L volume; 0.071 m^2^ cross sectional area) attached to a portable analyzer that determines CH_4_ concentrations in air using off-axis integrated cavity output spectroscopy (GLA132-GGA, ABB Measurement and Analytics; Ray et al. 2024). At the same time, we collected headspace samples for determination of dissolved CH_4_ concentrations. Using diffusive CH_4_ flux measurements and dissolved CH_4_ concentrations, we calculated gas transfer velocities (i.e., k_600_ values as:

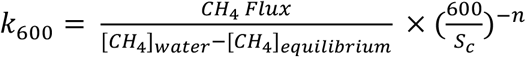

Where *S*_*c*_ and *n* are the Schmidt number for CH_4_ and the Schmidt number exponent, respectively.

### Water Column CH_4_ Bottle Incubations

We used light and dark bottle incubations to determine rates of CH_4_ production and consumption in surface and bottom waters of each pond. Immediately following the collection of dissolved CH_4_ samples, we filled 125 mL glass incubation bottles (one clear and one darkened with electrical tape, representative of ambient and low light conditions) and sealed them with gas tight rubber septa. These bottles were then suspended either 10 cm from the surface or 10 cm from the sediment-water interface. The following day, bottles were collected, de-capped, and sampled for temperature, dissolved oxygen, and dissolved CH_4_ concentrations as described previously. Rates of CH_4_ production or consumption were then calculated as the change in dissolved CH_4_ concentration over time, assuming starting conditions in each bottle matched the conditions in the pond when the bottles were filled.

### Potential Sediment CH_4_ Production

We collected sediments alongside microbial samples and stored them in acid-washed polyethylene bottles at −20 °C for later determination of potential sediment CH_4_ production using bench-top bottle incubations. Briefly, we added ∼ 5 g wet weight of thawed sediment to 125 mL glass incubation bottles equipped with optical O_2_ sensor spots (FireSting-GO2, PyroScience GMBH), added 70 mL of tap water that had been left out for > 24 hrs to allow for dissipation of any chlorine that might be present and to achieve atmospheric equilibrium, sealed the vials with gas tight rubber septa, and then over-pressured each vial with 15 mL of laboratory air. Two replicate bottles were used per pond. Headspace CH_4_ samples were collected on day 0, 2, 5, 6, and 14. Incubations were performed in the dark with room temperature ∼22 °C. To compare sediment CH_4_ production potential, we calculated rates of sediment CH_4_ production over time relative to the dry mass of sediment in each bottle. We note that this approach allows for comparison between ponds with and without FPV, but estimated CH_4_ production rates are unlikely to represent *in situ* conditions.

### Statistical Comparisons

All statistical analyses were performed using R statistical software and figures were made using the *ggplot2* package (Wickham 2016). We considered the results of statistical tests to be significant when p < 0.05.

We fit linear mixed-effects models (fixed: treatment, day of year; random: pond) to compare environmental and microbial metrics in FPV compared to Open ponds using *lme4* (Bates et al. 2015; Table S2). Rates of water column CH_4_ production or consumption in surface and bottom waters were compared using a mixed effects model of light or dark bottle and FPV presence with pond as a random effect (Table S3). Contrasts between treatments were made using least square means tests via *emmeans* (Lenth 2018) (Table S4).

Water temperature and oxygen were depth-averaged by pond and day. Diffusive CH_4_ fluxes, k600, and potential sediment CH_4_ production were compared using two-tailed t-tests (Tables S5&6).

Bray–Curtis dissimilarities were visualized with principal coordinates analysis (PCoA) and tested using PERMANOVA (adonis2), β-dispersion (betadisper), Mantel tests, and Procrustes analysis in *vegan* (Oksanen et al. 2018). Water column ASVs were analyzed as absolute abundances by scaling relative sequence abundances to total microbial cell counts (cells mL^-1^), whereas sediment ASVs were analyzed as relative abundances following rarefaction. Differentially abundant ASVs were identified with ANCOMBC-II (tax_level = “ASV”; prevalence >5%, fdr-adjusted q < 0.05, global test by FPV; Lin and Peddada 2024).

## Results

### FPV Ponds had Higher CH_4_ Concentrations at Depth

Dissolved CH_4_ concentrations were relatively stable over time (Fig. 1A & 1C). Surface CH_4_ did not differ between FPV (1.79 ± 4.01 μmol CH_4_ L^-1^) and control ponds (2.07 ± 3.19 μmol CH_4_ L^-1^; p = 0.208; Fig. 1B). However, bottom water CH_4_ was two orders of magnitude higher than in surface waters, and more than twice as high in FPV ponds (306.2 ± 419.0 μmol CH_4_ L^-1^) than controls (129.7 ± 45.60 μmol CH_4_ L^-1^; p = 0.003; Fig. 1D).

**Fig. 1:**
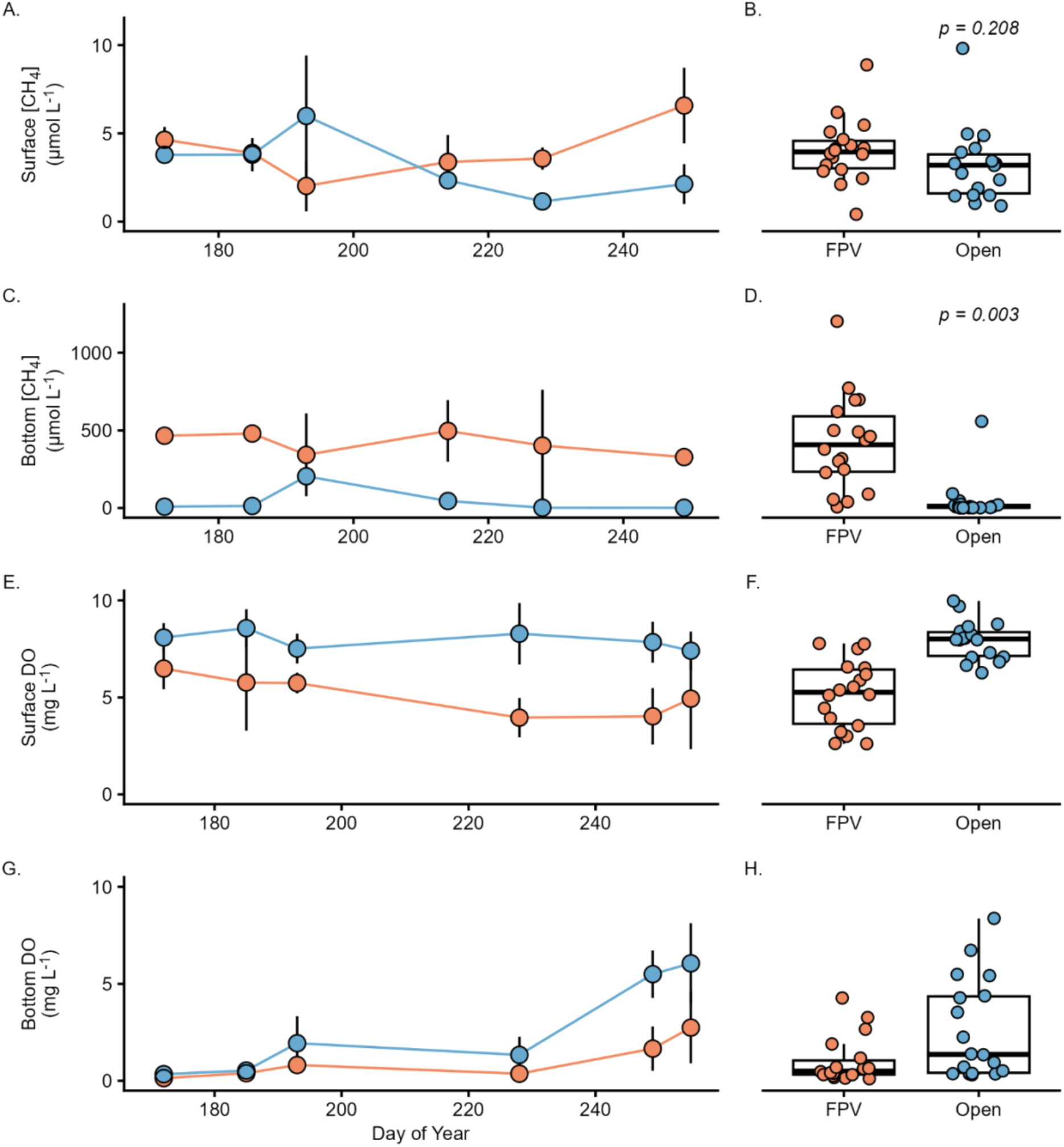
Dissolved methane concentrations (A-D) and dissolved oxygen (DO) concentrations (E-H) in the surface (A, B, E, F) and bottom water (C, D, G, H) of ponds with and without floating photovoltaic (FPV) arrays in summer 2024. In (A, C, E, G) points indicate the mean CH_4_ concentration from pond centers on a given date by treatment group and error bars indicate standard deviation. Colored lines connect the means. In panels (B, D, F, H) each point on the boxplot indicates a measured CH_4_ concentration in an individual pond based on whether FPV is present (FPV) or not (Open) and p-values indicate the result of a least-squares mean test of the treatment effect of the mixed model (Table S3).

### FPV Ponds were Colder and Less Oxygenated

Ponds with FPV were on average almost 4 °C colder across the water column (18.87 ± 1.69 °C; mean ± SD) than control ponds (22.78 ± 2.29 °C) during summer (p < 0.001; Fig. S1). On average, ponds could be considered stratified (i.e., density gradient > 0.29 kg m^-3^ m^-1^; Holgerson et al 2022), but we found no evidence that FPV changed pond mixing and stratification patterns, as mean density gradients in FPV ponds (0.76 ± 0.63 kg m^-3^ m^-1^) were similar to control ponds (0.62 ± 0.42 kg m^-3^ m^-1^; p = 0.180). While all ponds were generally undersaturated in DO, surface waters in control ponds were occasionally oversaturated, unlike FPV ponds, which remained undersaturated throughout the sampling period (Fig. S2; Table S7). Mean DO in FPV ponds (2.70 ± 0.98 mg L^1^) was half that of control ponds (5.42 ± 2.20 mg L^-1^; p < 0.001; Fig. 1 E-H & Fig. S3).

### FPV Ponds had Higher Water Column Methanotroph Abundance

Water column methanotroph abundance showed increasing divergence between FPV and Open ponds as the season progressed. In surface waters, methanotroph abundance was 3.7 times higher in FPV ponds (4.4 ± 3.7 x 10^5^ cells mL^-1^) than control ponds (1.2 ± 1.0 x 10^5^ cells mL^-1^; p = 0.021; Fig. 2A-B). Bottom waters displayed similar trends, with 2.8. times more methanotrophs in FPV ponds (Fig. 2E-H). Abundance of methanogens in the water column was orders of magnitude lower and there were no differences between treatments (Fig. 2C & D). Methanogens averaged ∼20% of the sediment community and were more abundant than methanotrophs (∼6%), but neither group differed between treatments (Fig. 2I-L).

**Fig. 2:**
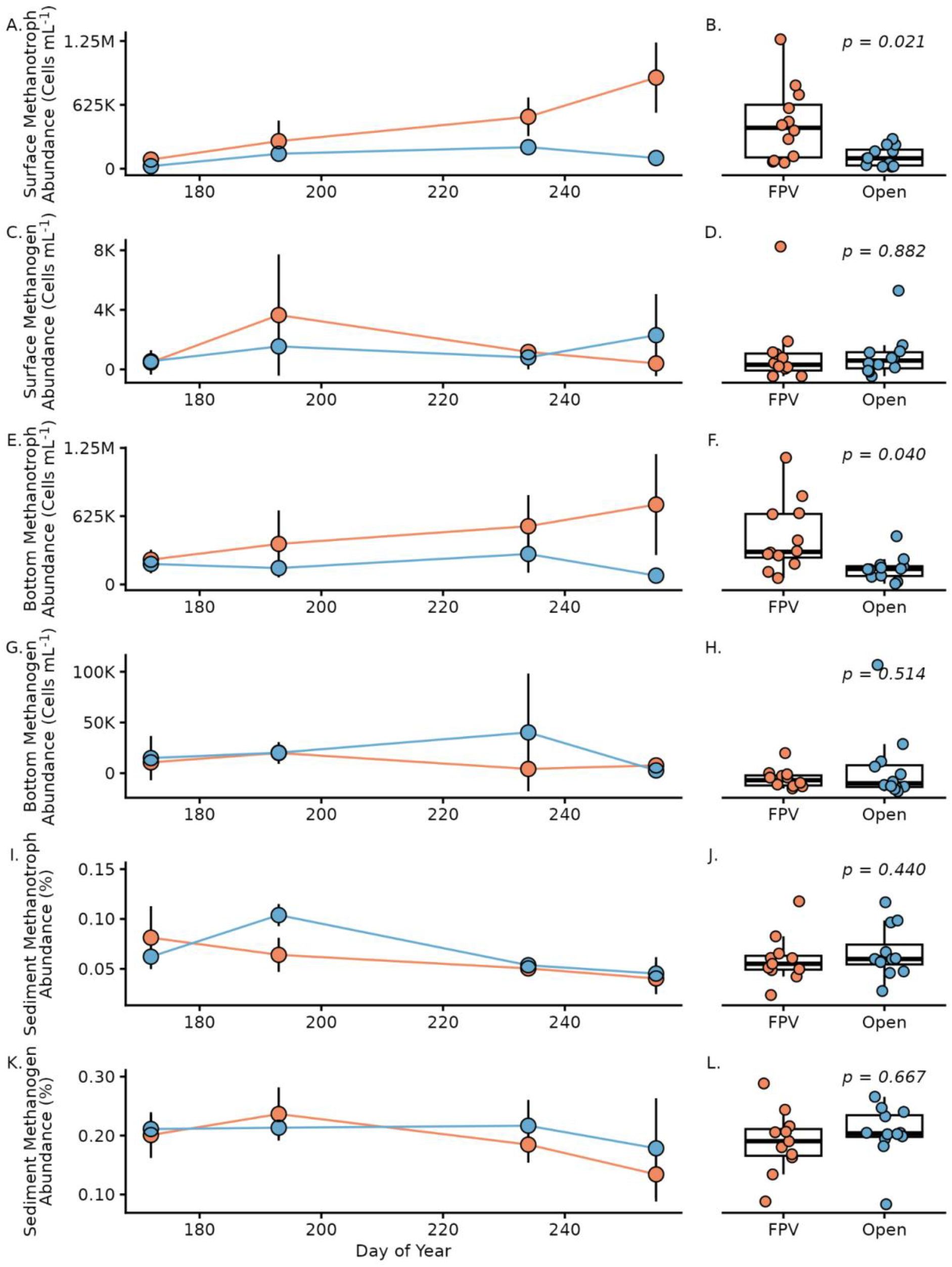
Total abundance (A-H) of methanotrophs and methanogens in surface waters and bottom waters and relative abundance (I-L) of methanotrophs and methanogens in sediments of FPV and Open ponds in summer 2024. Information as for Fig. 1. P-values reflect least-square means test result of the mixed model (Table S3).

### FPV Altered Methanotroph and Methanogen Community Structure

FPV altered methanotroph and methanogen community composition in both water and sediment (Fig. S4; Table S8), with FPV presence producing stronger between-group separation than pond identity (Table 1). Community composition varied with day of year (DOY), with significant FPV x DOY and pond x DOY interactions in both habitats. Spatial structuring by FPV presence and pond identity was more pronounced in sediments than in the water column. Water column community structure was tightly coupled with methanotroph composition rather than methanogens, indicated by strong correlations between water column community structure and methanotroph composition, supported by Mantel (r = 0.996, p < 0.001) and Procrustes (correlation = 0.999, m12 = 0.002, p < 0.001) analyses.

FPV altered seasonal trajectories in sediment methanogen communities, but not methanotroph communities. For both methanogens and methanotrophs, community composition was structured primarily by pond identity, with additional contributions from FPV and DOY, and for methanogens, a significant FPV x DOY interaction indicating FPV-specific seasonal shifts (Fig. S3A & B; Table S9).

FPV-associated shifts in water column methanotrophs were driven by a small number of Type 1 (hereafter gammaproteobacterial) methanotrophic ASVs identified by ANCOM-BC2. Four gammaproteobacterial methanotrophic ASVs within the order Methylococcales were 3-12x enriched in FPV ponds compared to controls. The strongest enrichment was observed for ASV_32 (*Methyloparacoccus*, family Methylococcaceae), which increased almost 12x under FPV. Additional enriched ASVs belonged to the family Methylomonadaceae, including ASV_13 (genus unresolved) and ASV_141 (*Methylobacter_C*), and ASV_44 (*Methylomonas*), which each increased ∼3-4x. Collectively, these individual taxa reached absolute abundances exceeding 300,000 cells mL^-1^ in FPV ponds. Four gammaproteobacterial methanotroph ASVs within the family Methylococcaceae decreased under FPV (∼3x lower abundance), including multiple ASVs classified as *Methyloterricola* (Fig. S4 & S5). These taxa occurred at much lower absolute abundances (Fig. S4 & S5), remaining one to two orders of magnitude less abundant than the FPV enriched methanotrophs (Fig. 4).

In sediments, FPV effects were not differentially abundant. However, there were treatment-specific seasonal trajectories across many of the most abundant sediment methanogen ASVs (Fig. S6), consistent with FPV x DOY interaction effects. These temporal shifts occurred across multiple ASVs within the genera *Methanothrix, Methanoregula, Methanosarcina, Methanobacterium* (Fig S6). Only a single ASV was enriched in the sediments, *Methanoperedens_A* (ASV_4603; an archaeal methanotroph), but its relative abundance was negligible.

### No Apparent Differences in CH_4_ Cycling Processes

We found no differences in diffusive water-air CH_4_ exchange or gas transfer velocities between ponds with and without FPV or between the center and edge of ponds (Table S5&10). On average, all ponds emitted CH_4_ to the atmosphere at a rate of 404.1 ± 896.6 μmol CH_4_ m^-2^ h^-1^. We also found no difference in rates of water column CH_4_ net-production or consumption between ponds with and without FPV (Table S3). Sediments from both Open and FPV ponds produced CH_4_, but we found no difference in potential CH_4_ production rates by sediments in ponds with FPV (5.43 ± 4.81 ppm CH_4_ g dry sediment^-1^ d^-1^) compared to those without (14.13 ± 10.92 ppm CH_4_ g dry sediment^-1^ d^-1^; p = 0.303; Table S6).

## Discussion

Following FPV deployment, bottom waters sustained elevated dissolved CH_4_ concentrations for a second year along with enhanced water column methanotroph abundances (Fig. 1D; Fig. 2). Surface water CH_4_ – and diffusive CH_4_ emissions – were similar between FPV and Open ponds. Together, these patterns suggest FPV-induced changes in redox conditions promote CH_4_ accumulation in bottom-waters, while microbially mediated CH_4_ oxidation dynamically constrains diffusive CH_4_ fluxes to the atmosphere, positioning water column methanotrophs as a potentially critical biofilter.

Specifically, we hypothesize that prolonged low-oxygen conditions in FPV ponds facilitate longer periods of CH_4_ accumulation in bottom waters overlying sediments. In response to this accumulation, water column methanotrophs reached bloom-like densities in late summer, exceeding 1,000,000 cells mL^-1^ and reaching abundances higher in FPV than Open ponds (Fig 2A). The coexistence of a persistently large CH_4_ pool and low, but not inhibitory, oxygen concentrations created conditions that selectively favored gammaproteobacterial methanotrophs later in the season, supporting elevated CH_4_ oxidation throughout the water column. By oxidizing CH_4_, methanotrophs serve as a fundamental biological sink that likely buffers diffusive CH_4_ emissions.

Beyond CH_4_ availability, FPV-driven hypoxia also restructured microbial composition in a lineage- and time-dependent manner. At the ASV level, closely related gammaproteobacterial methanotrophs within the order *Methylococcales* exhibited divergent responses to FPV across the season, with some lineages strongly enriched (Fig. 4 & S5) and others declining in FPV conditions (Fig S4 & S5). These taxa are adapted to low-oxygen environments and high CH_4_ availability (Li et al. 2024). Gammaproteobacterial methanotrophs utilize the ribulose monophosphate (RuMP) pathway for carbon assimilation, which is more energy efficient than the serine pathway used by alphaproteobacterial (Type II) methanotrophs and supports rapid growth under high CH_4_ availability (Knief, 2015). Therefore, FPV does not uniformly enhance water column methanotrophs, but instead acts as a selective filter that reshapes methanotroph community composition over time, promoting the growth of distinct *Methylococcales* genera at unique times of the season.

A similar FPV x DOY signal emerged within sediment methanogen communities. Although only relative abundance data were available for sediments and did not differ by FPV presence (Fig 2K-L), ASV-level time series revealed clear treatment-specific seasonal trajectories across many of the most abundant methanogen ASVs (Fig S6). These shifts included ASV-level changes in the timing and magnitude of mid-season peaks, often followed by earlier or steeper late-season declines relative to Open ponds (Fig S6). These same ASVs were not identified as differential abundance analysis, consistent with FPV effects operating primarily through time-dependent changes rather than a constant treatment effect as tested here. Because responses were distributed across many abundant ASVs rather than driven by a single taxon, sediment methanogen relative abundances did not differ between treatments (Fig. 2) despite strong compositional divergence (Fig 3 & Table S1&8).

**Fig. 3:**
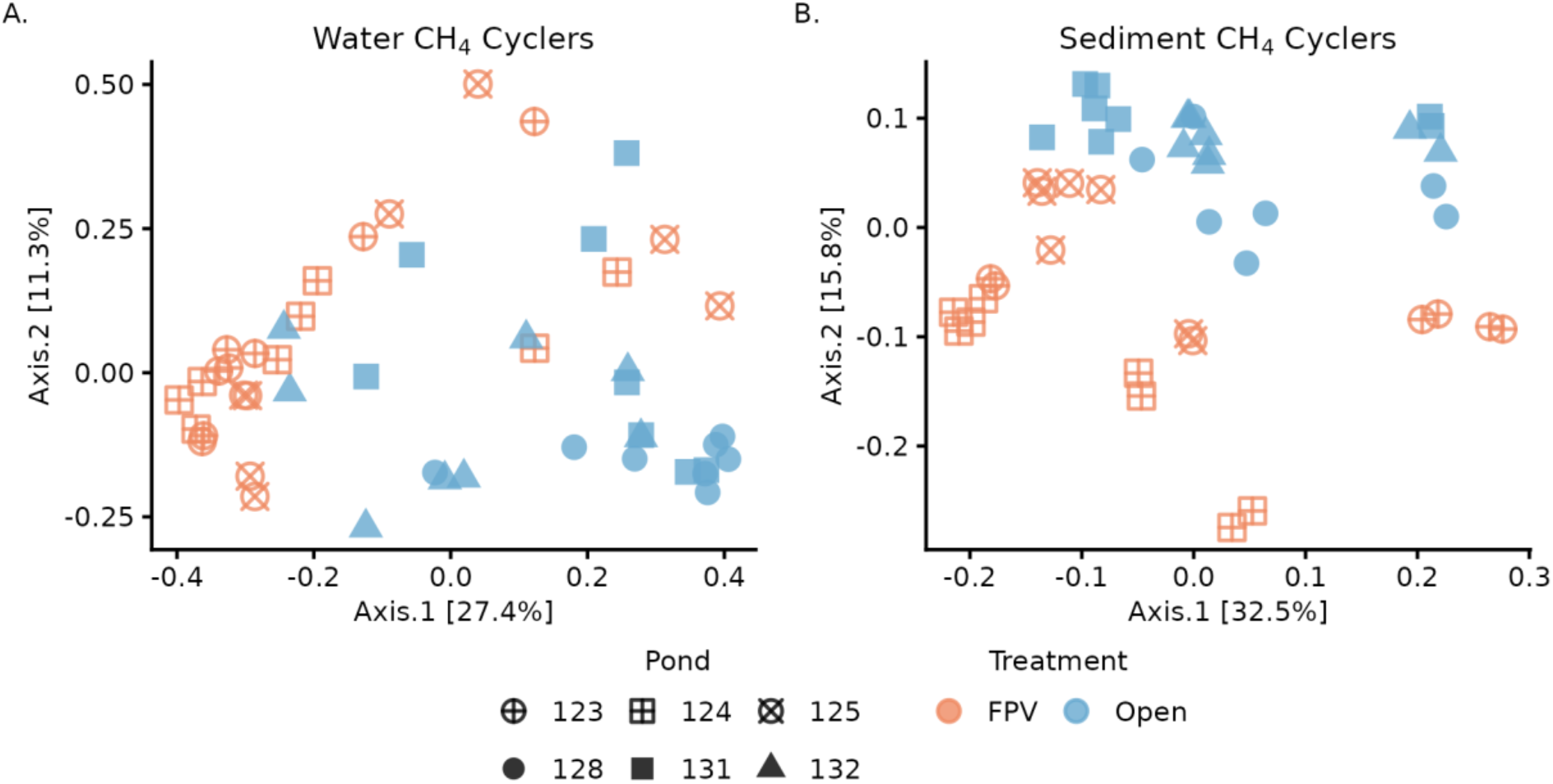
Bray–Curtis PCoA of combined methanogen and methanotroph communities in FPV and Open ponds. (A) Dissimilarity among water column samples was calculated using absolute cell abundances. (B) Sediment samples were rarefied to 20,826 reads to calculate dissimilarity based on relative abundance; replicates are shown separately. Shapes denote ponds.

**Fig. 4:**
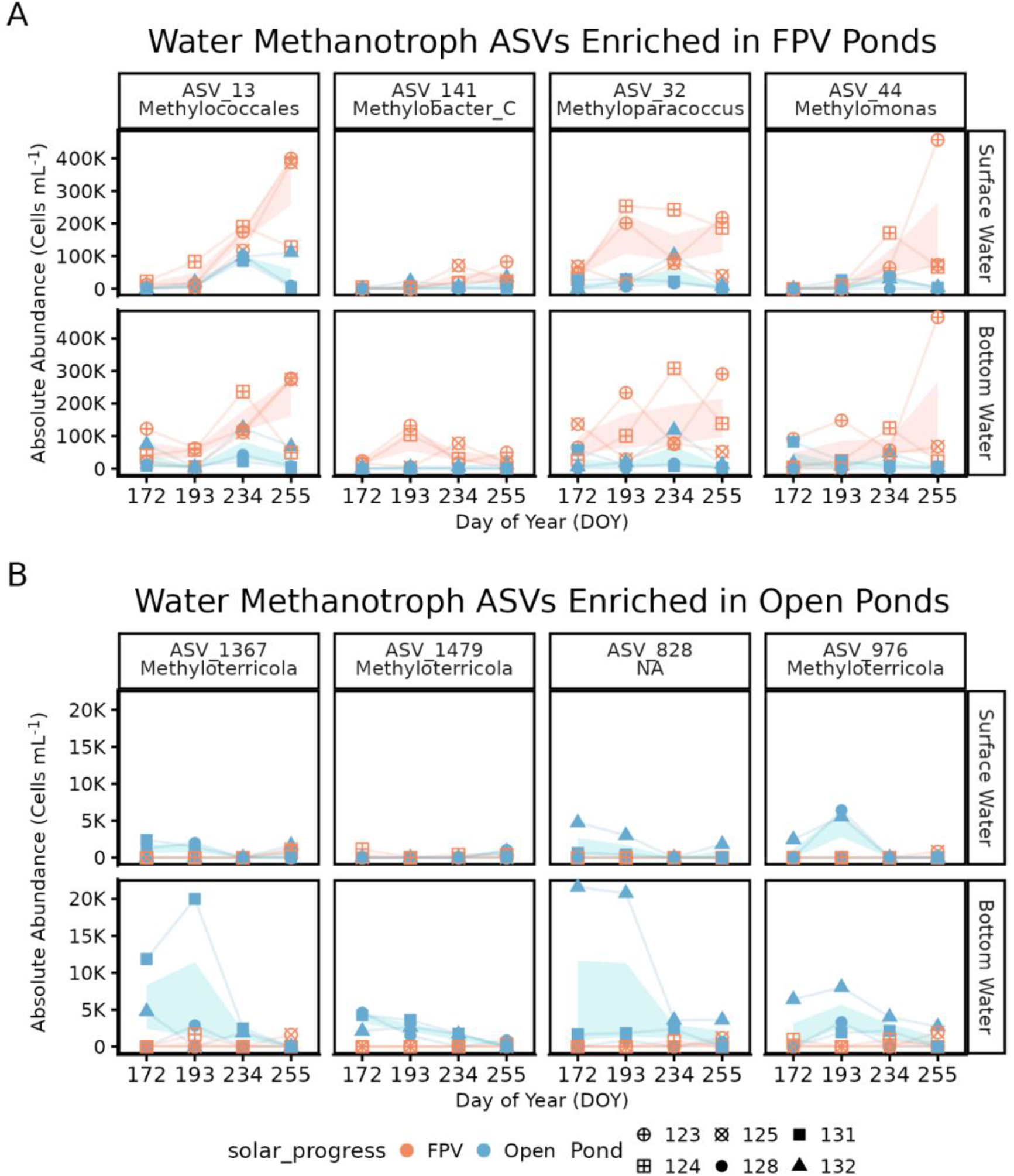
Methanotroph ASVs identified as differentially abundant between FPV and open ponds using ANCOM-BC2 are shown as absolute abundance (cells mL^−1^) across the sampling season. (A) Methanotroph ASVs that bloom under FPV infrastructure and (B) methanotroph ASVs enriched in open ponds. Panels are faceted by ASV and water-column depth (surface vs bottom). Points represent individual pond observations, lines connect repeated measurements through time within each pond–depth combination, and thick lines and shaded ribbons indicate treatment medians and interquartile ranges (25th–75th percentiles) at each date. FPV-enriched ASVs are dominated by members of the family Methylomonadaceae, with a single FPV-enriched Methylococcaceae ASV (*Methyloparacoccus*, ASV_32), whereas ASVs enriched in open ponds belong exclusively to Methylococcaceae. Taxonomic assignments are shown at the genus and species level, except ASV_13, which is classified within Methylomonadaceae, and ASV_828, which is classified within Methylococcaceae. For visualization, two low-abundance FPV-enriched ASVs (ASV_2028, ASV_346) and one open-pond ASV with minimal separation in absolute abundance (ASV_1019) were excluded.

**Fig. 5:**
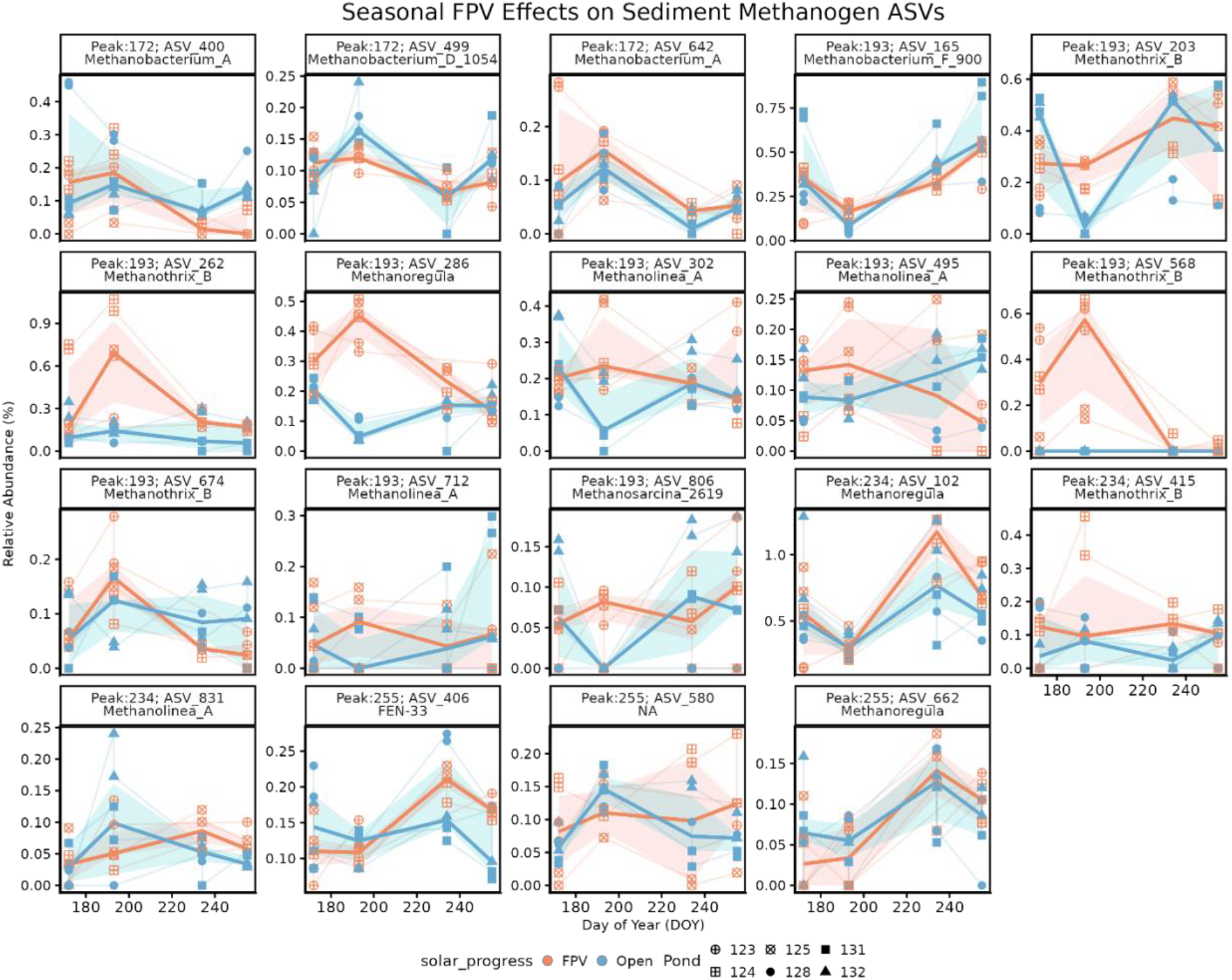
Sediment methanogen ASVs showing strong, time-dependent divergence between FPV and open ponds are shown as relative abundance (%) across the sampling season. To identify FPV-associated seasonal patterns not captured by a constant treatment effect, methanogen ASVs were screened using a manual, seasonally explicit approach focused on abundant taxa (mean relative abundance > 0.05%). FPV–Open separation was quantified at each sampling date using median abundances, and ASVs were retained if peak FPV enrichment was supported across multiple sampling dates and/or multiple FPV ponds (see Supplemental Methods). Panels are faceted by ASV and annotated with the day of year (DOY) at which maximum FPV enrichment occurred. Points represent individual pond observations, thin lines connect repeated measurements within ponds, and thick lines and shaded ribbons indicate treatment medians and interquartile ranges (25th–75th percentiles) at each date. ASVs span multiple methanogenic phyla and classes, including genera within hydrogenotrophic (*Methanobacterium*), acetoclastic (*Methanothrix*), and methylotrophic (*Methanoregula, Methanosarcina*) groups.

In late summer, methanotrophs surpassed 1,000,000 cells mL^-1^ in FPV surface and bottom waters, a density rarely reported in any aquatic system. This bloom may reflect the metabolic advantage of gammaproteobacterial methanotrophs under CH_4_-rich, low-O_2_ conditions, as observed in freshwater lakes where gammaproteobacterial methanotrophs proliferate under hypoxia and anoxia (Reis et al. 2020; Reis et al. 2024). And while we did not measure light availability, shading from FPV floats and panels may also have enhanced methanotrophic activity (Murase & Sugimoto 2005; Thottathil et al. 2018).

Our results also demonstrate that the effect of FPV on pond biogeochemistry and CH_4_ cycling varies over time. Immediately following FPV deployment in 2023, FPV ponds had lower oxygen concentrations and increased surface and bottom water CH_4_ concentrations (Ray et al. 2024). By contrast, in the second year (2024), surface CH_4_ concentrations no longer differed between treatments, and methanotroph abundance was substantially higher in FPV surface waters. These patterns suggest a time-lagged ecological response, where sustained hypoxia promotes bottom-water CH_4_ accumulation, followed by methanotroph enrichment that buffers CH_4_ flux near the surface. Quantifying how CH_4_ ebullition responds to FPV deployment across seasons and years remains a key open question for understanding total emission dynamics.

Identifying strategies for low emission energy production is critical for combating climate change. As FPV expansion accelerates, understanding how this technology interacts with aquatic CH_4_ dynamics is important for ensuring sustainability. Our study provides mechanistic evidence that FPV deployment can restructure microbial communities in ways that influence carbon cycling, offering a foundation for evaluating the long-term ecosystem-level impacts of continued FPV proliferation.

## Acknowledgements

This work was supported by an Atkinson Academic Venture Fund to SMG, MAH, MLS, awards 441993/2023-0 and 200781/2024-3 from National Council for Scientific and Technological Development (CNPq) to SJC, and startup funds from the University of Delaware made available to NER. We thank Benj Sterrett for helping to maintain access to the ponds, Autumn Newman, Augustus Pendleton, Sophia Richter and Kailyn Hanke for assistance with field and laboratory work, and Jera Jansen, Caitlin Davis, Mônica Antunes-Ulyssea, Sheena Dwyer-McNulty, Trifosa Simamora, Tim Boycott, and Dave Grodsky for helping construct the floating solar arrays. Any use of trade, firm, or product names is for descriptive purposes only and does not imply endorsement by the U.S. Government.

## Author Contribution Statement

NER designed the study, provided supplies, performed field sampling, laboratory analyses, and data analysis, and wrote the first draft of the manuscript. SA performed field sampling, laboratory analyses, data analysis, and edited the manuscript. SMG secured funding, constructed the floating solar arrays, performed field sampling, and edited the manuscript. AC, SJC, and MT performed field sampling and edited the manuscript. MAH secured funding, performed field sampling, and edited the manuscript. MLS designed the study, provided supplies, data analysis, secured funding, and edited the manuscript.

## Scientific Significance Statement

Producing energy using floating photovoltaic (FPV) powerplants offers an opportunity to produce renewable energy, spare land, and reduce evaporation from ponds, lakes, and reservoirs. However, FPV deployment in these ecosystems is associated with colder temperatures, less oxygen availability, and changes in carbon cycling processes. Specifically, experimental evidence demonstrates an increase in concentrations of methane – a potent greenhouse gas – in ponds following FPV deployment. In this study, we investigate how microbial communities associated with aquatic methane cycling differ between ponds with and without FPV. We show that FPV deployment increases bottom-water methane concentrations and triggers dense blooms of methane-oxidizing bacteria. These results provide the first evidence that microbial communities respond strongly to engineered shading and may help buffer greenhouse gas emissions in solar-covered waters.

## Data and Code Availability

All raw and processed data for this project are publicly available. The code used for statistical comparisons, generating figures, and processing microbial community data are available on GitHub at https://github.com/MarschmiLab/Ray_LO_Letters_FPV_Methane. Biogeochemical data and summaries of methane cycling microbe abundances is available for download via the Figshare Repository (https://doi.org/10.6084/m9.figshare.29614025.v1. The raw, compressed 16S rRNA gene sequencing fastq files are available in the NCBI Sequence Read Archive (SRA) under the BioProject accession number PRJNA1417770 (https://www.ncbi.nlm.nih.gov/bioproject/1417770). All flow cytometry data are available from Zenodo (https://zenodo.org/records/18462017).

