## supplemental for "Higher methanotroph abundance and bottom-water methane in ponds with floating photovoltaic arrays"

*This information product has been peer reviewed and approved for publication as a preprint by the U.S. Geological Survey.*

#### Supplementary Material

### Higher methanotroph abundance and bottom-water methane in ponds with floating photovoltaic arrays

Any use of trade, firm, or product names is for descriptive purposes only and does not imply endorsement by the U.S. Government.

Nicholas E. Ray[1], Sophia Aredas[2], Steven M. Grodsky[3], Ash Canino[4], Simone J. Cardoso[5], Meredith A. Holgerson[6], Meredith Theus[6], Marian L. Schmidt[2]

[1] School of Marine Science & Policy, University of Delaware, USA
[2] Department of Microbiology, Cornell University, USA
[3] U.S. Geological Survey, New York Cooperative Fish and Wildlife Research Unit, Department of Natural Resources and the Environment, Cornell University, USA
[4] Department of Natural Resources and Environment, Cornell University, USA
[5] Department of Zoology, Institute of Biological Sciences, Universidade Federal de Juiz de Fora, Brazil
[6] Department of Ecology and Evolutionary Biology, Cornell University, US

#### Supplemental Methods

##### Water Column Microbial Cell Abundances

Water samples were preserved for flow cytometry by adding 1 µL of 25% glutaraldehyde to 1 mL of 20 µm pre-filtered pond water, shaking, incubating for 10 minutes in the dark at room temperature, and freezing at -80 °C. Fixed samples were diluted to 15-20X, stained with SYBR Green I, and run in triplicate on an Attune NxT Flow Cytometer. DNA-positive cell counts were analyzed using the *flowCore* (Ellis et al. 2025) and *ggcyto* (Van et al. 2018) packages in R (R Core Team 2014).

##### Testing for Water Column Stratification

We tested if the water column was stratified for each pond on each sampling date by comparing calculated density gradients between ponds with and without FPV. We used temperature measurements in the surface and bottom waters, corrected for the distance between those samples. Water density at a given temperature was calculated as:

$$\rho(T) = \frac{\rho_{H2O}}{1 + \beta(T - T_0)} \times \frac{1000\ kg\ m^{-3}}{1\ kg\ L^{-1}}$$

Where ρ(T) is the density of water at a given temperature (kg $m^{-3}$), $\rho_{H2O}$ is the nominal density of water (assuming 1 kg $L^{-1}$ for freshwater), β is the volumetric temperature expansion coefficient (0.0002 $°C^{-1}$), T is the temperature of the water (C), and $T_0$ is standard temperature (20 °C).

We then calculated the density gradient between the surface and bottom water temperature (kg $m^{-3}$ $m^{-1}$) for each pond on each sampling day, and compared density gradients between ponds with and without FPV by constructing a mixed effect model with FPV presence and day of year as fixed effects and pond as a random effect, using least square means to compare density gradients between ponds with and without FPV.

R Core Team. 2014. R: A language and environment for statistical computing.

**Table S1**: Methanogen and Methanotroph Taxonomic Classification

| Order | Family | Genus | Methane Cycler |
|---|---|---|---|
| *Methanocellales* | *Methanocellaceae* | *Methanocella_A* | Methanogen |
| *Methanomicrobiales* | *Methanomicrobiaceae* | | Methanogen |
| *Methanomicrobiales* | *Methanospirillaceae_2121* | *Methanolinea_A*<br>*Methanolinea_B*<br>*Methanoregula*<br>*UBA467*<br>*UBA288* | Methanogen |
| *Methanomicrobiales* | *Methanospirillaceae_2125* | *Methanospirillum* | Methanogen |
| *Methanosarcinales_A_2632* | *Methanosarcinaceae* | *Methanosarcina_2619*<br>*Methanomethylovorans* | Methanogen |
| *Methanotrichales* | *Methanotrichaceae* | *Methanothrix_B* | Methanogen |
| *Methanobacteriales* | *Methanobacteriaceae* | *Methanobacterium_A*<br>*Methanobacterium_B_963*<br>*Methanobacterium_C*<br>*Methanobacterium_D_1054*<br>*Methanobacterium_F_896*<br>*Methanobacterium_F_900*<br>*Methanobrevibacter_A*<br>*Methanobrevibacter_D*<br>*Methanosphaera* | Methanogen |
| *Methanofastidiosales* | *Methanofastidiosaceae* | *Methanofastidiosum* | Methanogen |
| *Methanomassiliicoccales* | *Methanomassiliicoccaceae* | *Methanomassiliicoccus_A_1624* | Methanogen |
| *Methanomassiliicoccales* | *Methanomethylophilaceae* | | Methanogen |

| | | | |
|---|---|---|---|
| *Methanomassiliicoccales* | *UBA472* | *FEN-33* | Methanogen |
| *Methanomethylicales* | *Methanomethylicaceae* | *Methanomethylicus* | Methanogen |
| *Methanosarcinales_A_2632* | *Methanoperedenaceae* | *Methanoperedens_A* | Methanotroph |
| *Methylomirabilales* | *2-02-FULL-66-22* | *2-02-FULL-66-22* | Methanotroph |
| *Rhizobiales_505101* | *Beijerinckiaceae* | *Methylocystis*<br>*Methylosinus* | Methanotroph |
| *Methylococcales* | *Methylococcaceae* | *Methylocaldum*<br>*Methylococcus*<br>*Methylomagnum*<br>*Methyloparacoccus*<br>*Methyloterricola*<br>*Methylotetracoccus*<br>*UBA6136* | Methanotroph |
| *Methylococcales* | *Methylomonadaceae* | *Crenothrix*<br>*Methylobacter_C_601048*<br>*Methylobacter_C_601751*<br>*Methyloglobulus*<br>*Methylomonas*<br>*Methylosoma*<br>*Methylovulum*<br>*UBA4132* | Methanotroph |

**Table S2:** Summary of model information for mixed models to describe temperature, dissolved oxygen, surface and bottom water dissolved methane ($CH_4$) concentration, methanotroph abundance, and methanogen abundance, relative abundance of sediment methanogens and methanotrophs, and density gradients in ponds with and without floating photovoltaics (FPV). All models include FPV presence or absence and day of year as fixed effects and the pond ID as a random effect. For each fixed effect, the estimated effect size is listed ± standard error of the effect, with the associated t-value below in parentheses.

| Response | Observations (n) | Intercept | FPV Presence | Day of Year |
|---|---|---|---|---|
| Temperature | 36 | 24.9 ± 2.06<br>(12.1) | 3.91 ± 0.60<br>(6.46) | -0.03 ± 0.01<br>(-3.00) |
| Dissolved Oxygen<br>(% Saturation) | 36 | -19.2 ± 18.5<br>(-1.03) | 32.2 ± 5.43<br>(5.92) | 0.23 ± 0.08<br>(2.72) |
| Dissolved Oxygen<br>(mg $L^{-1}$) | 36 | -3.45 ± 1.61<br>(-2.14) | 2.72 ± 0.53<br>(5.13) | 0.03 ± 0.01<br>(3.93) |
| Surface Water [$CH_4$] | 36 | 5.61 ± 2.60<br>(2.16) | -0.82 ± 0.65<br>(-1.26) | -0.01 ± 0.01<br>(-0.62) |
| Bottom Water [$CH_4$] | 36 | 631 ± 289<br>(2.18) | -373 ± 126<br>(-2.96) | -1.02 ± 1.33<br>(-0.77) |
| Surface water methanotroph abundance | 24 | -626754 ± 287292<br>(-2.18) | -318325 ± 86085<br>(-3.70) | 4997 ± 1316<br>(3.80) |
| Surface water methanogen abundance | 24 | -1332 ± 2632<br>(0.51) | -125 ± 789<br>(-0.16) | 0.44 ± 12.05<br>(0.04) |
| Bottom water methanotroph abundance | 24 | -94652 ± 324453<br>(-0.29) | -291441 ± 97220<br>(-3.00) | 2623 ± 1485<br>(1.77) |
| Bottom water methanogen abundance | 24 | 24108 ± 28147<br>(0.40) | 8895 ± 12441<br>(0.51) | -63.13 ± 125<br>(-0.62) |
| Sediment methanotroph abundance | 23 | 0.15 ± 0.03<br>(5.51) | 0.007 ± 0.008<br>(0.86) | -0.000 ± 0.000<br>(-3.42) |
| Sediment methanogen abundance | 23 | 0.32 ± 0.05<br>(5.83) | 0.014 ± 0.030<br>(0.46) | -0.001 ± 0.000<br>(-2.47) |
| Density Gradient<br>(kg $m^{-3}$ $m^{-1}$) | 36 | 3.53 ± 0.37<br>(9.54) | -0.145 ± 0.108<br>(-1.34) | -0.013 ± (0.002)<br>(-7.65) |

**Table S3:** Summary of model information for mixed models to describe bottle incubations to determine rates of water column dissolved methane ($CH_4$) production or consumption in ponds with and without floating photovoltaics (FPV). All models include FPV presence or absence and if the bottle was light or dark as fixed effects and the pond ID as a random effect. For each fixed effect, the estimated effect size is listed ± standard error of the effect, with the associated t-value below in parentheses.

| Response | Observations (n) | Intercept | FPV Presence/Absence | Light/Dark Bottle |
|---|---|---|---|---|
| Surface $CH_4$ Flux | 12 | -0.09 ± 0.07 (-1.27) | -0.08 ± 0.10 (-0.82) | 0.08 ± 0.03 (3.01) |
| Bottom $CH_4$ Flux | 12 | -1.69 ± 4.14 (-0.41) | 2.11 ± 5.83 (0.36) | -0.66 ± 0.88 (-0.75) |

**Table S4:** Results of least-square means comparisons between ponds with and without floating photovoltaics. Models for which comparisons are made are described in Tables S1 and S2. For bottle incubations to assess water column methane transformations, a positive value indicates net methane production, while a negative value indicates net methane consumption.

| Comparison | Estimate ± SE | z-ratio | p-value |
|---|---|---|---|
| Temperature (°C) | -3.91 ± 0.60 | -6.46 | < 0.001 |
| Dissolved Oxygen (% Saturation) | -32.3 ± 5.43 | -5.92 | < 0.001 |
| Dissolved Oxygen (mg $L^{-1}$) | -2.72 ± 0.53 | -5.13 | < 0.001 |
| Surface Water [$CH_4$] | 0.82 ± 0.65 | 1.26 | 0.208 |
| Bottom Water [$CH_4$] | 373 ± 126 | 2.96 | 0.003 |
| Surface Water $CH_4$ Production/Consumption | 0.08 ± 0.10 | 0.82 | 0.413 |
| Bottom $CH_4$ Production/Consumption | -2.11 ± 5.83 | -0.36 | 0.718 |
| Surface water methanotroph abundance | 318325 ± 86100 | 3.70 | 0.021 |
| Surface water methanogen abundance | 125 ± 789 | 0.158 | 0.882 |
| Bottom water methanotroph abundance | 291441 ± 97200 | 3.00 | 0.040 |
| Bottom water methanogen abundance | -8895 ± 12400 | -0.72 | 0.514 |
| Sediment methanotroph abundance | -0.007 ± 0.008 | -0.858 | 0.440 |
| Sediment methanogen abundance | -0.014 ± 0.030 | -0.46 | 0.667 |
| Density Gradient | 0.145 ± 0.108 | 1.40 | 0.180 |

**Table S5:** Results of Welch two-sample t-tests comparing diffusive methane fluxes and gas exchange velocities (k600) between the center and edges of ponds with and without floating photovoltaic arrays (FPV). We first compared pond locations across treatments, and then between locations and within treatments.

| Comparisons Between Ponds with and Without FPV | | | | |
|---|---|---|---|---|
| Measurement | Pond Location | df | t-value | p-value |
| Diffusive $CH_4$ Flux | Center | 2.00 | -1.03 | 0.413 |
| Diffusive $CH_4$ Flux | Edge | 2.00 | -0.97 | 0.436 |
| k600 | Center | 2.18 | -0.75 | 0.525 |
| k600 | Edge | 2.00 | -1.03 | 0.412 |
| Comparisons between pond Center and Edge within the same pond type | | | | |
| Measurement | FPV Present/Absent | df | t-value | p-value |
| Diffusive $CH_4$ Flux | Present | 2.28 | -1.67 | 0.221 |
| Diffusive $CH_4$ Flux | Absent | 3.34 | 0.34 | 0.751 |
| k600 | Present | 2.03 | 1.17 | 0.362 |
| k600 | Absent | 3.42 | -0.31 | 0.774 |

**Table S6:** Results of Welch two-sample t-test comparing potential rates of sediment $CH_4$ production between ponds with and without floating photovoltaic arrays.

| Measurement | df | t-value | p-value |
|---|---|---|---|
| Sediment $CH_4$ Flux | 2.75 | -1.26 | 0.302 |

**Table S7:** Summary of mean ± standard deviation of measured concentrations and rates in ponds with FPV and control ponds without FPV. For potential sediment $CH_4$ production and water column $CH_4$ "flux", positive values indicate net-$CH_4$ production while negative values indicated net-consumption.

| Measurement | Control Ponds | FPV Ponds |
|---|---|---|
| Temperature (°) | 22.78 ± 2.29 | 18.87 ± 1.69 |
| Dissolved Oxygen (% Saturation) | 62.34 ± 22.79 | 30.18 ± 10.56 |
| Dissolved Oxygen (mg $L^{-1}$) | 5.42 ± 2.19 | 2.70 ± 0.98 |
| Surface Water [$CH_4$] (µmol $L^{-1}$) | 3.19 ± 2.07 | 4.01 ± 1.79 |
| Bottom Water [$CH_4$] (µmol $L^{-1}$) | 45.6 ± 129.7 | 419 ± 306 |
| Surface water methanotroph abundance | | |
| Surface water methanogen abundance | | |
| Bottom water methanotroph abundance | | |
| Bottom water methanogen abundance | | |
| Sediment methanotroph abundance | | |
| Sediment methanogen abundance | | |
| Density Gradient (kg $m^{-3}$ $m^{-1}$) | 0.62 ± 0.42 | 0.76 ± 0.63 |
| Potential Sediment $CH_4$ Production (ppm $CH_4$ g dry $sediment^{-1}$ $d^{-1}$) | 14.1 ± 10.9 | 5.43 ± 4.81 |
| Surface Water Column $CH_4$ "Flux" (µmol $CH_4$ $L^{-1}$ $hr^{-1}$) | -0.14 ± 0.15 | -0.05 ± 0.08 |
| Bottom Water Column $CH_4$ "Flux" (µmol $CH_4$ $L^{-1}$ $hr^{-1}$) | 0.09 ± 1.58 | -2.02 ± 9.03 |

**Table S8:** FPV treatment, pond, and day of year (DOY) significantly influence the combined methanogen and methanotroph community composition in the water column and sediments. Bray-Curtis dissimilarities were calculated using absolute abundances for water (Fig. 3A) and scaled relative abundances for sediment (Fig. 3B). PERMANOVA tested effects of FPV, pond, and DOY and their interactions on community composition, with the pseudo-F and $R^2$ representing group separation variance explained, respectively. β-dispersion tested group variance. Residual $R^2$ shown for each habitat.

| Habitat Type | Variable | PERMANOVA | | | | β-Dispersion | | Figure |
|---|---|---|---|---|---|---|---|---|
| | | $R^2$ | Pseudo-F | p-value | Residuals | F | p-value | |
| Water | FPV | 0.128 | 9.42 | 0.001 | 0.489 | 10.23 | 0.002 | 3A |
| Water | Pond | 0.155 | 2.85 | 0.001 | | 1.20 | 0.305 | 3A |
| Water | DOY | 0.086 | 6.35 | 0.001 | | 5.26 | 0.005 | 3A |
| Water | FPV:DOY | 0.059 | 4.34 | 0.001 | | | | 3A |
| Water | Pond:DOY | 0.083 | 1.52 | 0.041 | | | | 3A |
| Sediment | FPV | 0.135 | 12.02 | 0.001 | 0.361 | 1.51 | 0.216 | 3B |
| Sediment | Pond | 0.285 | 6.32 | 0.001 | | 0.66 | 0.68 | 3B |
| Sediment | DOY | 0.107 | 9.48 | 0.001 | | 1.73 | 0.172 | 3B |
| Sediment | FPV:DOY | 0.030 | 2.67 | 0.018 | | | | 3B |
| Sediment | Pond:DOY | 0.082 | 1.81 | 0.011 | | | | 3B |

**Table S9:** FPV, pond, and day of year (DOY) structure sediment methanogen and methanotroph communities with distinct drivers and effect strengths. PERMANOVA and β-dispersion results for sediment methanogen and methanotroph communities. Bray-Curtis dissimilarities were calculated from rarefied relative abundances. FPV, pond, and DOY were modeled as main effects and interactions. $R^2$ indicates the proportion of variance explained; pseudo-F and p-values assess compositional differences. β-dispersion statistics test group variability. Residual $R^2$ is reported for each $CH_4$ group. Figures S3A and S3B show corresponding ordinations.

| Habitat Type | $CH_4$ Group | Variable | PERMANOVA | | | | β-Dispersion | | Figure |
|---|---|---|---|---|---|---|---|---|---|
| | | | $R^2$ | Pseudo-F | p-value | Residuals | F | p-value | |
| Sediment | Methanogens | FPV | 0.135 | 12.4 | 0.001 | 0.349 | 2.92 | 0.105 | S3A |
| Sediment | Methanogens | Pond | 0.305 | 7.01 | 0.001 | | 0.809 | 0.557 | S3A |
| Sediment | Methanogens | DOY | 0.098 | 9.01 | 0.001 | | 1.03 | 0.411 | S3A |
| Sediment | Methanogens | FPV:DOY | 0.033 | 3.06 | 0.012 | | | | S3A |
| Sediment | Methanogens | Pond:DOY | 0.079 | 1.82 | 0.016 | | | | S3A |
| Sediment | Methanotrophs | FPV | 0.131 | 10.5 | 0.001 | 0.398 | 0.004 | 0.937 | S3B |
| Sediment | Methanotrophs | Pond | 0.231 | 4.63 | 0.001 | | 0.723 | 0.604 | S3B |
| Sediment | Methanotrophs | DOY | 0.133 | 10.7 | 0.001 | | 4.13 | 0.009 | S3B |
| Sediment | Methanotrophs | FPV:DOY | 0.020 | 1.59 | NS | | | | S3B |
| Sediment | Methanotrophs | Pond:DOY | 0.088 | 1.76 | 0.018 | | | | S3B |

**Table S10:** Summary of diffusive CH4 fluxes, dissolved CH4 concentrations, and calculated $k_{600}$ values for the edges and centers of ponds with and without FPV. Values reported as mean ± standard deviation.

| Measurement | Location | Control | FPV |
|---|---|---|---|
| Diffusive $CH_4$ Flux (µmol $m^{-2}$ $hr^{-1}$) | Center | 954 ± 1565 | 26.7 ± 5.6 |
| Diffusive $CH_4$ Flux (µmol $m^{-2}$ $hr^{-1}$) | Edge | 588 ± 969 | 47.8 ± 21.1 |
| Dissolved $CH_4$ Concentration (µmol $L^{-1}$) | Center | 5.99 ± 3.43 | 2.02 ± 1.44 |
| Dissolved $CH_4$ Concentration (µmol $L^{-1}$) | Edge | 3.08 ± 1.03 | 6.18 ± 2.47 |
| k600 (cm $h^{-1}$) | Center | 0.40 ± 0.62 | 0.12 ± 0.13 |
| k600 (cm $h^{-1}$) | Edge | 0.60 ± 0.96 | 0.03 ± 0.01 |

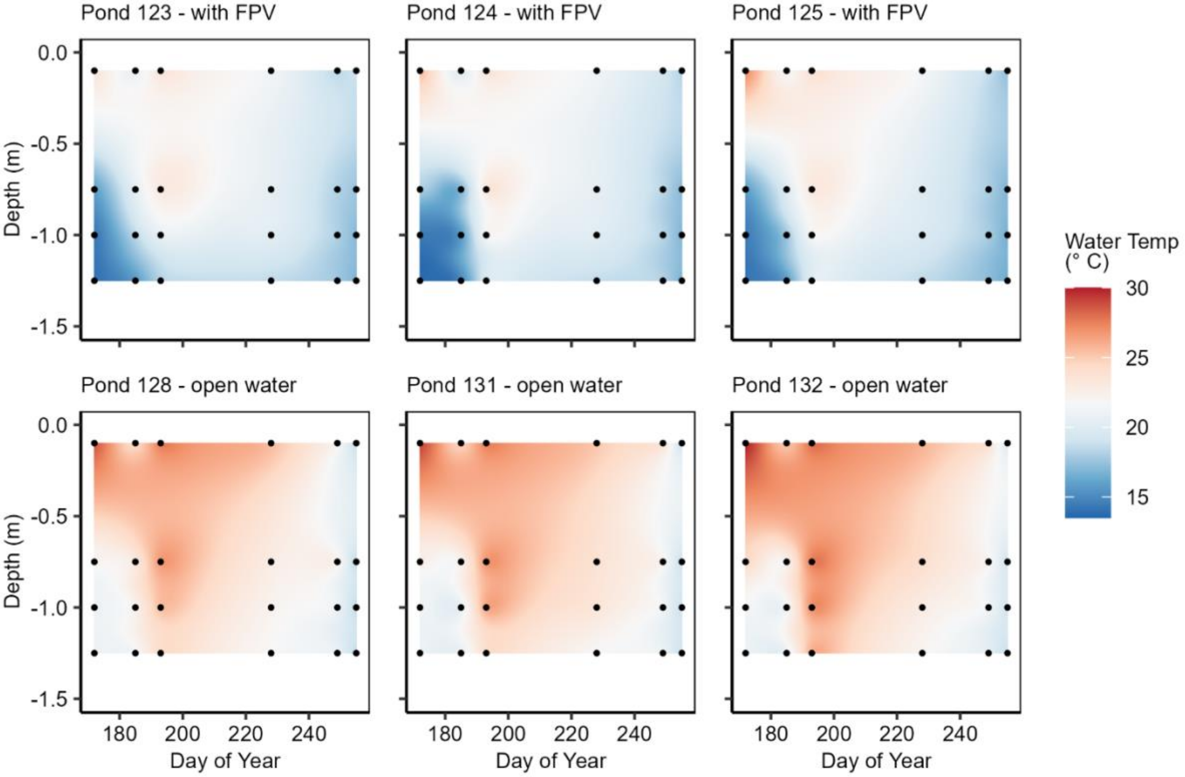

**Fig. S1:** Temperature profiles throughout summer 2024 in ponds with floating photovoltaic (FPV) arrays (top row) and ponds without FPV arrays (bottom row). Points indicate measurements and colors are interpolated based on those measurements.

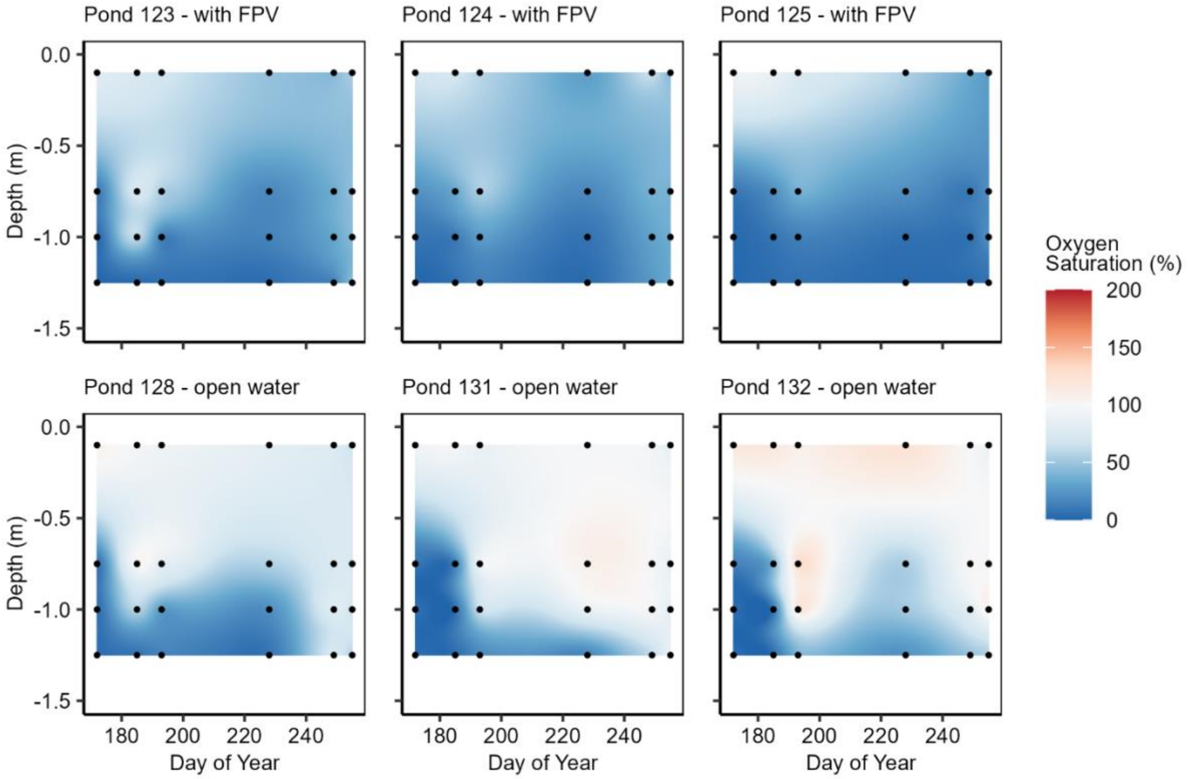

**Fig. S2:** Dissolved oxygen (percent saturation) profiles throughout summer 2024 in ponds with floating photovoltaic (FPV) arrays (top row) and ponds without FPV arrays (bottom row). Points indicate measurements and colors are interpolated based on those measurements.

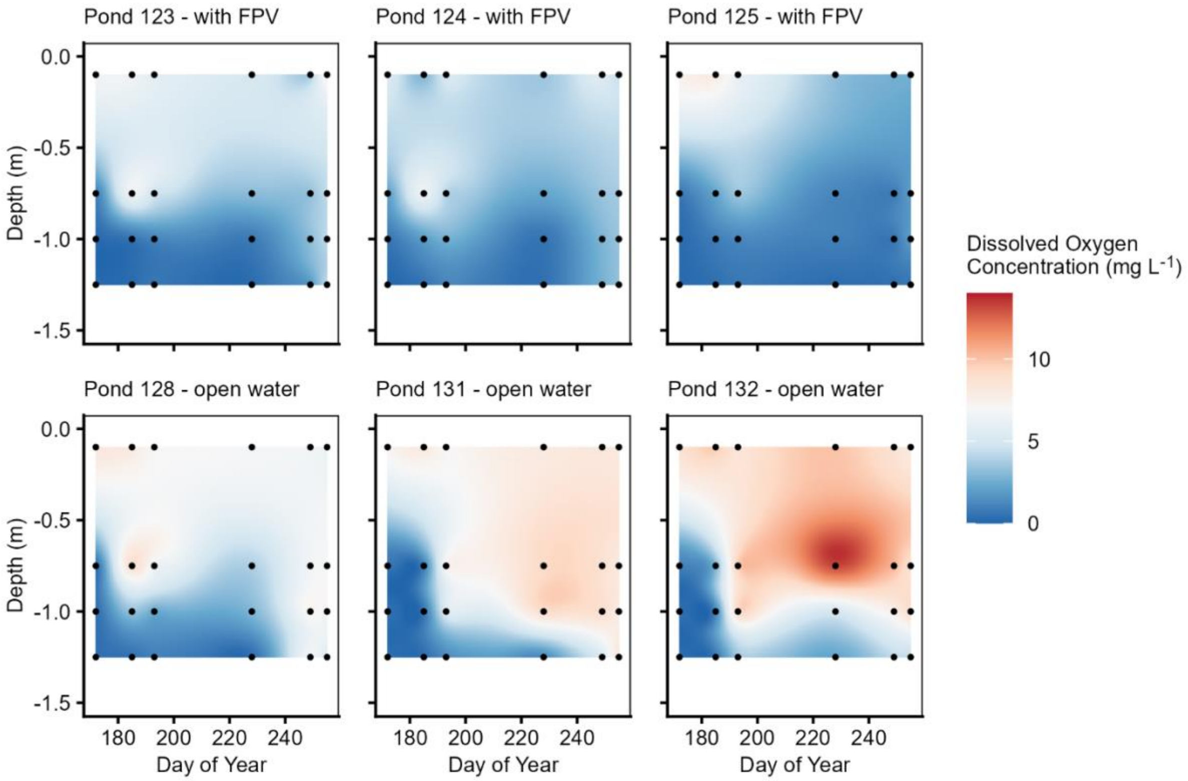

**Fig. S3:** Dissolved oxygen (mg $L^{-1}$) profiles throughout summer 2024 in ponds with floating photovoltaic (FPV) arrays (top row) and ponds without FPV arrays (bottom row). Points indicate measurements and colors are interpolated based on those measurements.

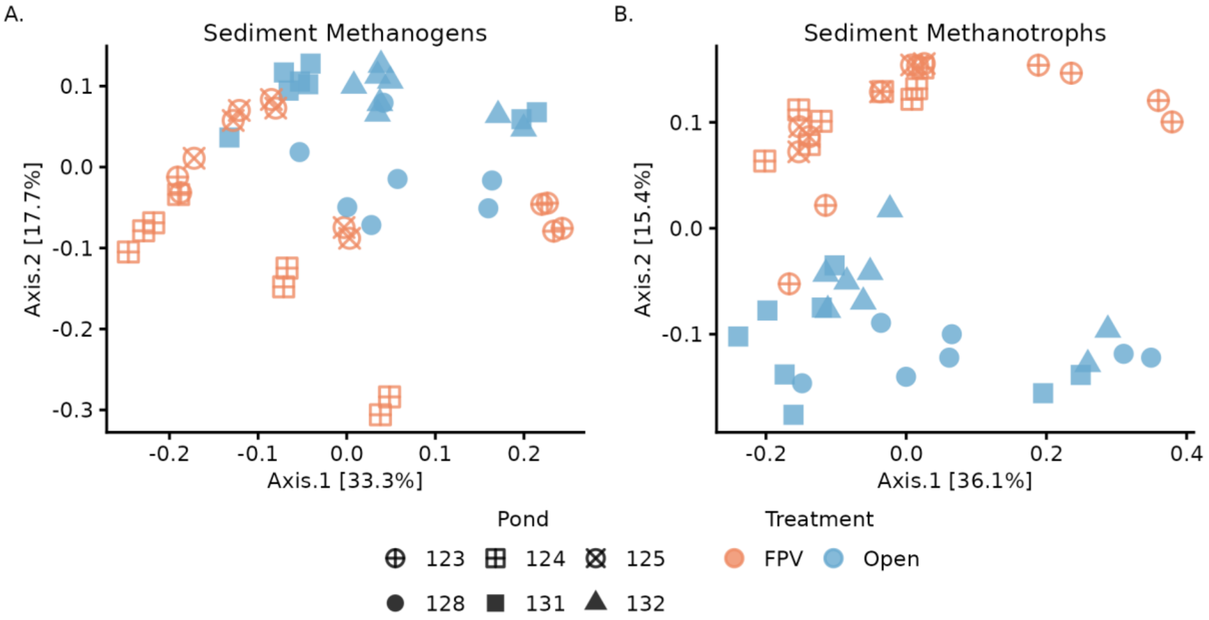

**Fig. S4**: Bray-Curtis PCoA of sediment (A) methanogenic and (B) methanotrophic communities within FPV and Open ponds; Sediment replicates and are shown separately.

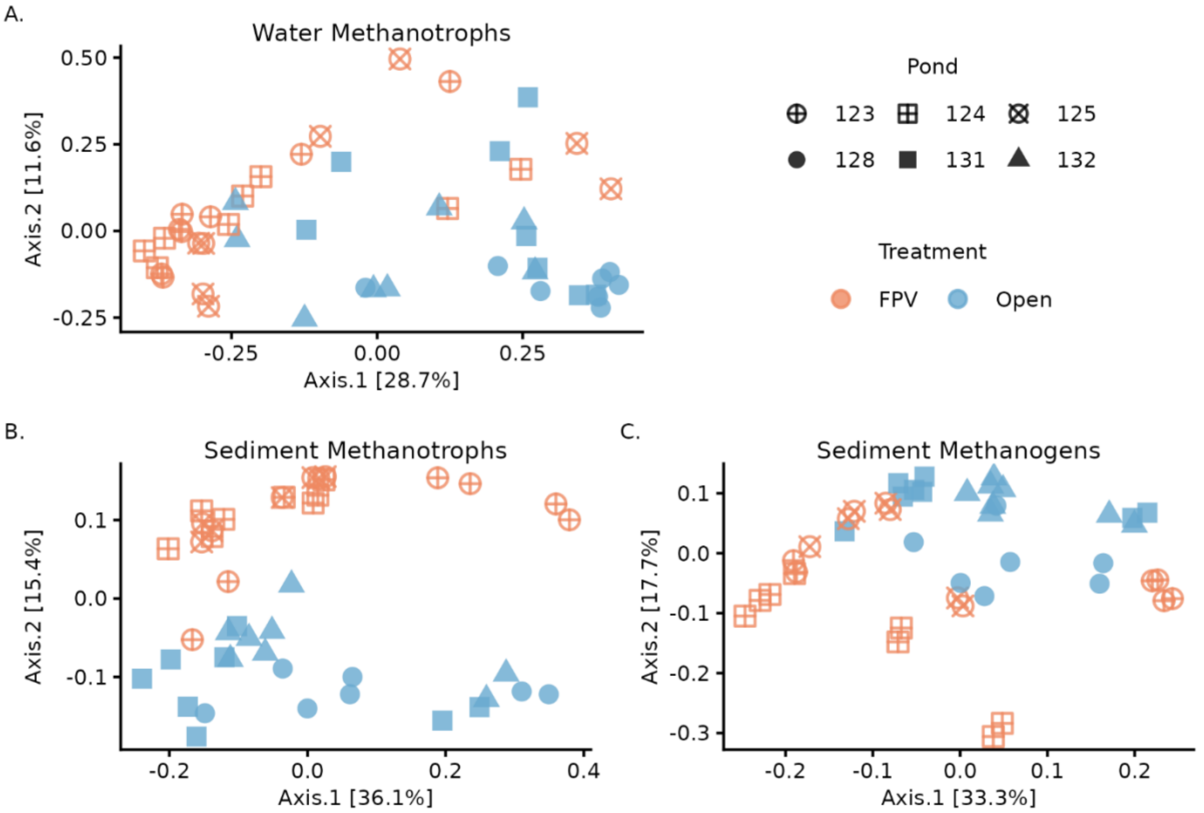

**Fig. S5**: Bray-Curtis PCoA of (A) water methanotrophs, sediment (B) methanotrophs and (C) methanogen communities within ponds with FPV ("FPV") and ponds without FPV ("Open"); Sediment replicates and are shown separately.

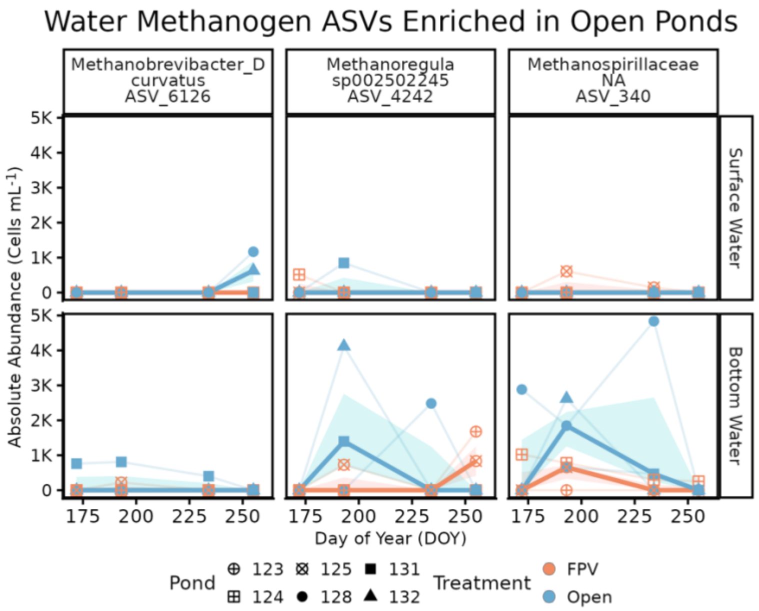

**Fig. S6.** Time-averaged differential abundance of water-column methanogen ASVs under non-FPV pond conditions. Absolute abundances (cells $mL^{-1}$) of water-column methanogen ASVs identified as enriched in open ponds using time-averaged ANCOM-BC2 analyses are shown across the sampling season.

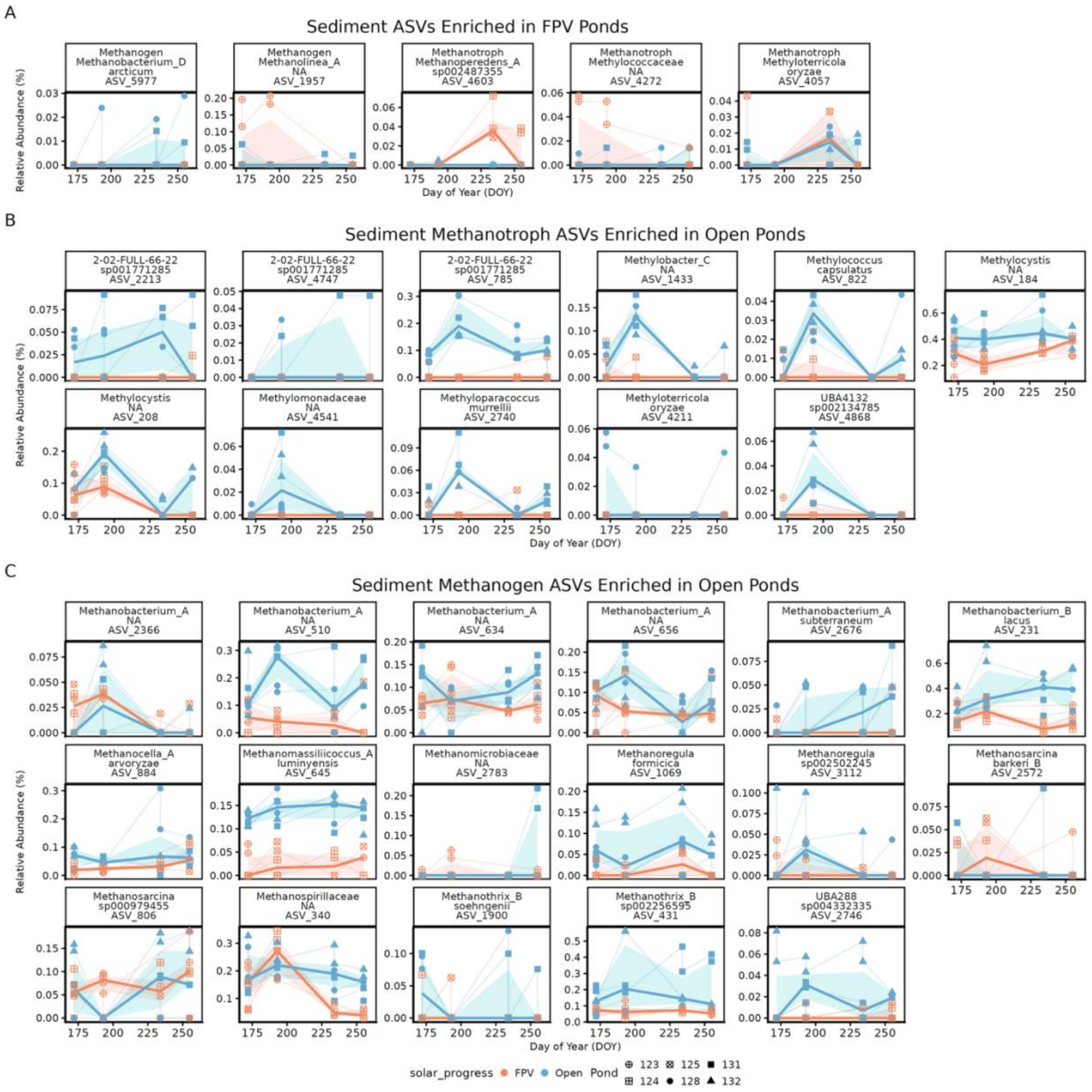

**Fig. S7.** Time-averaged differential abundance of sediment methane-cycling ASVs under FPV and non-FPV pond ("Open") conditions. Sediment methane-cycling ASVs identified as differentially abundant using time-averaged ANCOM-BC2 analyses. (A) ASVs enriched in FPV ponds, (B) methanotroph ASVs enriched in open ponds, and (C) methanogen ASVs enriched in open ponds. Relative abundance (%) of each ASV is shown across day of year (DOY). ASVs lacking genus-level taxonomic resolution are labeled at the family level in facet titles (ASV_340, ASV_2783, ASV_4541, ASV_2366, ASV_634).

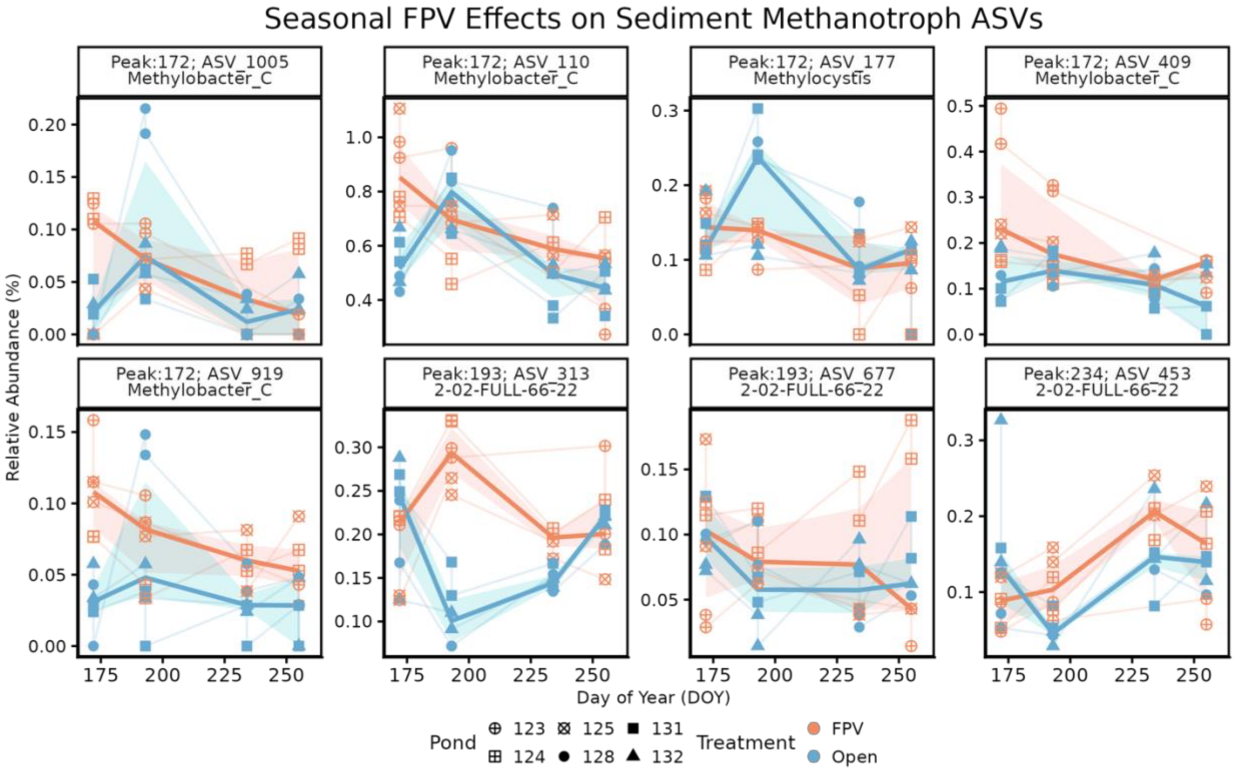

**Fig. S8.** Seasonally explicit enrichment of sediment methanotroph ASVs under FPV conditions. Relative abundance (%) of selected sediment methanotroph ASVs across the sampling season, highlighting ASVs with peak FPV-associated enrichment identified using a seasonally explicit (time-resolved) screening approach rather than time-averaged differential abundance testing. Taxa include Gammaproteobacteria (Type I; *Methylobacter_C*), Alphaproteobacteria (Type II; *Methylocystis*), and anaerobic nitrite-dependent methanotrophs (Type III; *Methylomirabilales*, 2-02-FULL-66-22). Community-level analyses did not detect a significant FPV × time effect for sediment methanotrophs.
